# Deep Generative Models for Detecting Differential Expression in Single Cells

**DOI:** 10.1101/794289

**Authors:** Pierre Boyeau, Romain Lopez, Jeffrey Regier, Adam Gayoso, Michael I. Jordan, Nir Yosef

**Author notes:** Corresponding author. (N.Y.).

## Abstract

Detecting differentially expressed genes is important for characterizing subpopulations of cells. However, in scRNA-seq data, nuisance variation due to technical factors like sequencing depth and RNA capture efficiency obscures the underlying biological signal. First, we show that deep generative models, which combined Bayesian statistics and deep neural networks, better estimate the log-fold-change in gene expression levels between subpopulations of cells. Second, we use Bayesian decision theory to detect differentially expressed genes while controlling the false discovery rate. Our experiments on simulated and real datasets show that our approach out-performs state-of-the-art DE frameworks. Finally, we introduce a technique for improving the posterior approximation, and show that it also improves differential expression performance.

## 1 Introduction

Detecting differential gene expression (DE) is important for characterizing subpopulations of cells profiled by single-cell RNA sequencing (scRNA-seq) [1]. Technical factors like sequencing depth and RNA capture efficiency obscure the underlying biological signal in scRNA-seq measurements, making DE difficult [2]. Many methods have been developed to specifically handle the complexities of scRNA-seq [3, 4, 5], each with distinct data-generating distributions and hypothesis formulations, but they do not consistently outperform DESeq2 [6] and edgeR [7], which were designed for bulk expression data [8]. This might be attributable to the lack of flexibility of the underlying statistical models (generalized linear models fit to each gene separately, with limited information shared between cells).

Conversely, a new class of methods uses deep neural networks as nonlinear functions to model conditional distributions. In scVI [9] for example, the mean expression of each gene is a nonlinear function of a cell-specific latent variable that represents biological state. Hypothesis testing in scVI consists of posterior sampling and Bayes factor computation. As parameters are shared between all cells, this approach is agnostic to the partition of the data. It is particularly convenient since it avoids fitting a different model for each test (e.g., given two clusters to compare) and allows a better quantification of gene expression levels uncertainty. However, the scVI hypothesis formulation might lead to detection of biologically irrelevant genes, without further thresholding on the effect size (namely, log fold change, LFC). Also, the mean-field approximate posterior that it used to fit the model may poorly represent the true posterior, limiting the model’s accuracy.

Our main contribution is to employ deep generative models for LFC estimation and differential expression by extending the scVI framework in order to address the limitations of existing methods. We use Bayesian decision theory [10] to explicitly control for effect-size via LFC (Section 2). Our experiments on simulated and real datasets show that our approach outperforms state-of-the-art DE frameworks (Section 3). Finally, we show that an alternative posterior approximation scheme that may better capture uncertainty further improves differential expression performance. These results suggest that deep generative models (DGM) are valuable tools for uncertainty quantification in gene expression data.

## 2 Methods

### 2.1 Background

A scRNA-seq experiment produces observations *x*_*ng*_, which represent in a cell *n* the number of mRNA transcripts mapping to gene *g*. In scVI, a cell’s gene expression is generated in the following way. First, a low-dimensional latent variable *z*_*n*_ representing a cell’s biological state is sampled for a standard multivariate normal distribution. This sample of *z*_*n*_ is mapped by a neural network to another latent variable *h*_*ng*_, representing the underlying expression level for gene *g*. Next, a scalar latent variable *l*_*n*_ representing the cell’s sequencing depth and size is sampled from a log normal distribution. The observation model for *x*_*ng*_ is a zero-inflated negative binomial distribution, where the mean of the negative binomial component is equal to the product *h*_*ng*_*l*_*n*_. We assume that latent parameters of *z*_*n*_ can be mapped to a cell type or cell state via a deterministic mapping (e.g., a clustering algorithm).

While we focus on the scVI model in this work, our framework for differential expression can be applied to any generative model parametrized by *θ* of the form

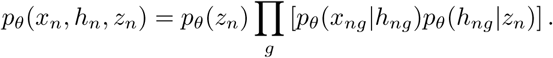

### 2.2 Effect-size control in differential expression

Given two cells *a* and *b* we represent their difference in expressing a gene *g* using a latent variable 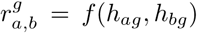, where the function *f* is designed to capture some biologically meaningful change in magnitude between its inputs. We define *f* to be the LFC, log_2_ *h*_*ag*_ *−* log_2_ *h*_*bg*_, which is a popular measure of effect-size for expression measurements.

Let 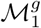 (resp. 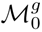) be the model for which gene *g* is (resp. is not) differentially expressed. It is expected that biologically meaningful shifts in gene expression occurs when the LFC is (in absolute value) higher than a threshold *δ*, defined by the practitioner [6]. We consequently rely on random variables 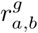 to define formally differential expression by

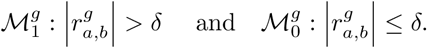

By design, this differential expression formulation filters out gene that might be significantly DE for *δ′ < δ* (i.e., of low magnitude) but are likely not to be interesting in practice.

Let *K*_0_ be the cost of a false negative and *K*_1_ the cost of a false positive. If 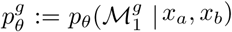 is the posterior probability of gene *g* being DE at a sufficient LFC, Bayesian decision theory [10] shows that the optimal decision rule consists of calling genes which satisfy

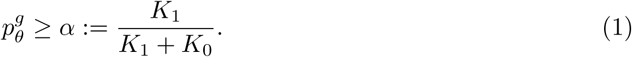

Another potentially interesting design would be to control the false discovery rate, as in [11]. In this manuscript, however, we focus on estimating 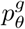.

### 2.3 Log-fold change estimation

The posterior LFC distribution can be obtained via marginalization of latent variables:

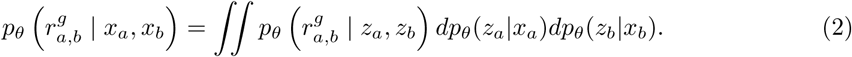

Consequently, 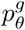 can be approximated using Monte Carlo given access to the posterior distribution *p*_*θ*_(*z*_*n*_ *| x*_*n*_). This definition can be generalized to arbitrary pairs of cell sets *A* = (*a*_1_, …, *a*_*n*_) and *B* = (*b*_1_, …, *b*_*m*_) via the following distribution:

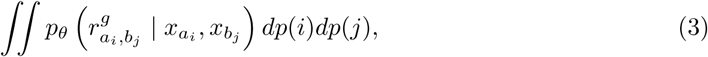

where *i ∼ 𝒰* (*a*_1_, …, *a*_*m*_) and *j ∼ 𝒰* (*b*_1_, …, *b*_*m*_). This is tantamount to sampling from the aggregate posteriors 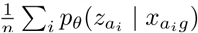 and 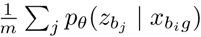 instead of the individual posteriors in Equation 2. When *A* and *B* correspond to two groups of cells of distinct types, the aggregated LFC defined in Equation 3 converges to a oracle cell-type-specific LFC with enough samples (not shown in this manuscript). The aggregation noted in Equation 3 might be suboptimal in the presence of outliers or cell type mislabeling, though we leave the treatment of such cases as future work.

### 2.4 Variational approximation

We fit *p*_*θ*_ and learn an approximate posterior distribution *q*_*ϕ*_(*z*_*n*_ *| x*_*n*_) using the AEVB framework [12]. The variational posterior *q*_*ϕ*_(*z*_*n*_ *| x*_*n*_) can be used as a proxy for the real posterior *p*_*θ*_(*z*_*n*_ *| x*_*n*_) in Equation 2. The quality of the variational distribution *q*_*ϕ*_ can play an important role for this estimation. In vanilla scVI [9] and in AEVB [12], the Gaussian mean and diagonal covariance of the variational distribution is parameterized via encoder networks (referred to as naive mean-field, MF). However, a mean-field factorization of the variational distribution for *z*_*n*_ is known to underestimate the posterior variance [13]. Such behavior is expected to affect the coverage of LFC estimates as well as the downstream decision rule for differential expression. To underscore this, we will compare MF against inverse autoregressive flows (IAF) [14], which provides richer posterior approximations, and therefore learns better model with improved uncertainty quantification.

## 3 Results

We are interested in two tasks: LFC estimation (with analysis of the mean predictions as well as uncertainties) and differential expression detection (i.e., estimating *p*_*θ*_). For both tasks, we apply our framework using scVI with two choices of variational approximations to the posterior: MF and the more expressive IAF.

Furthermore, we compare these models to MAST [3], DESeq2 [6] and edgeR [7] using the implementation of [15], which correct p-values for multiple hypothesis testing using false discovery rate control (at significance threshold 0.05). In DESeq2, a Wald test based on shrunken LFC estimates is used for differential expression. Similarly to our approach, DESeq2 formulates composite null hypothesis in which the LFC absolute value is below a certain threshold *δ*. We use *δ* = 0.5 for both DESeq2 and our method.

In the tables, starred values represent significantly better results compared to DESeq2, edgeR and MAST at level 0.05. Bold values denote significantly better results among our methods (MF and IAF).

### 3.1 Datasets

We assess performance on both synthetic and real datasets. The first synthetic dataset (PL) consists of 8,000 cells of two cell-types, with cells in each type generated from a Poisson log-normal distribution. Ground-truth LFCs between the two cell-types are computed as the average log-ratios of generated Poisson rates. We use a dense covariance matrix to model gene-gene interactions for the log-normal distribution. The DE genes in this synthetic dataset are those for which the true LFC magnitude is above 0.5.

We also employ SymSim [16] to study the robustness of predictions to scRNA-seq technical noise on a dataset of 10,000 cells and 1,000 genes corresponding to two batches (UMI and non-UMI) of five cell types. SymSim does not provide exact normalized LFCs that we can take as ground truth, but they can be approximated from the kinetic parameters given by the simulator [16].

Finally, we consider a peripheral blood mono-nuclear cells (PBMC) dataset composed of 12,039 cells and 3,346 genes from two batches, as described in [9]. Additional bulk data allows to provide inter cell-types microarray LFC estimates that we will consider to be ground-truth.

### 3.2 LFC estimation

In this section, we consider the LFC estimation task and show that our pipeline has competitive performance in both real and synthetic datasets.

We first compare scVI LFC mean estimates on the PL synthetic dataset to the other methods as a function of the query size, which corresponds to the number of cells of each cell-type taken as input for LFC estimation by the different algorithms. The bigger the query size, the closer we can expect the predicted LFC to match the ground-truth. scVI provides significantly better estimates (Table 1), especially for small query sizes.

**Table 1:**
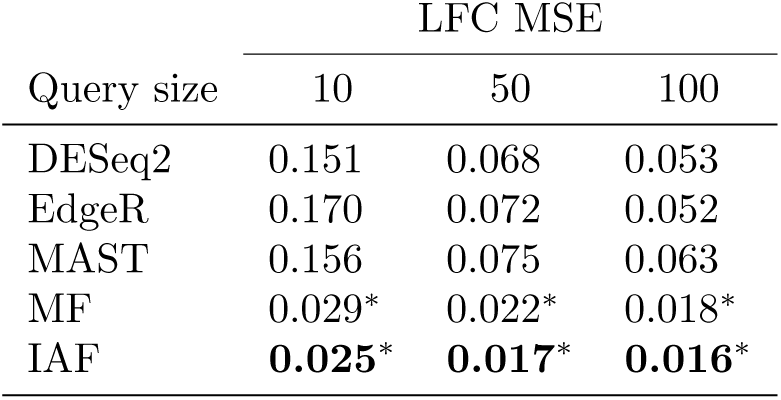
LFC mean squared error for different population sizes. Each algorithm’s predictions are run for ten different cells-samplings and for five weight initializations for scVI-based methods.

Such behavior is expected as scVI fits a single model for the entire dataset while the other methods are fit only on the cells being compared. When the query size is small, other techniques show instability and important errors that scVI does not suffer from. Note that IAF predictions outperform MF, suggesting that it allows to fit a better generative model.

As our LFC inference protocol is Bayesian, the study of the LFC posterior distributions can provide helpful uncertainty information about the log-fold changes estimates. Figure 1 suggests that DGMs can successfully estimate LFC on scRNA-seq data and that credible intervals can give valuable insight into the estimations. That being said, the models uncertainty estimates differ slightly as IAF credible intervals are often shorter than MF. Hence, The question that arises consists in determining which variational distribution models uncertainty the most faithfully. To further investigate this point, we compare MF and IAF calibration errors as defined in [17]. Let *p*_1_, … *p*_*m*_ be given confidence levels. Let 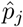 be the fraction of genes for which the real LFC in contained is the *p*_*j*_-credible intervals of some model. The calibration error 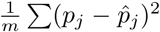 will reflect the quality of the estimated uncertainty of the model. We compute the calibration errors of posterior LFCs of the different models for 20 random subsets of cells for confidence levels 10%, 20%, 30%, 40%. IAF and MF mean calibration errors respectively are 0.0019 and 0.0031 (Mann-Whitney U-test p-value *<* 0.05). This improved modelling of uncertainty show that LFCs credible intervals are better estimated by more complex variational distributions.

**Figure 1:**
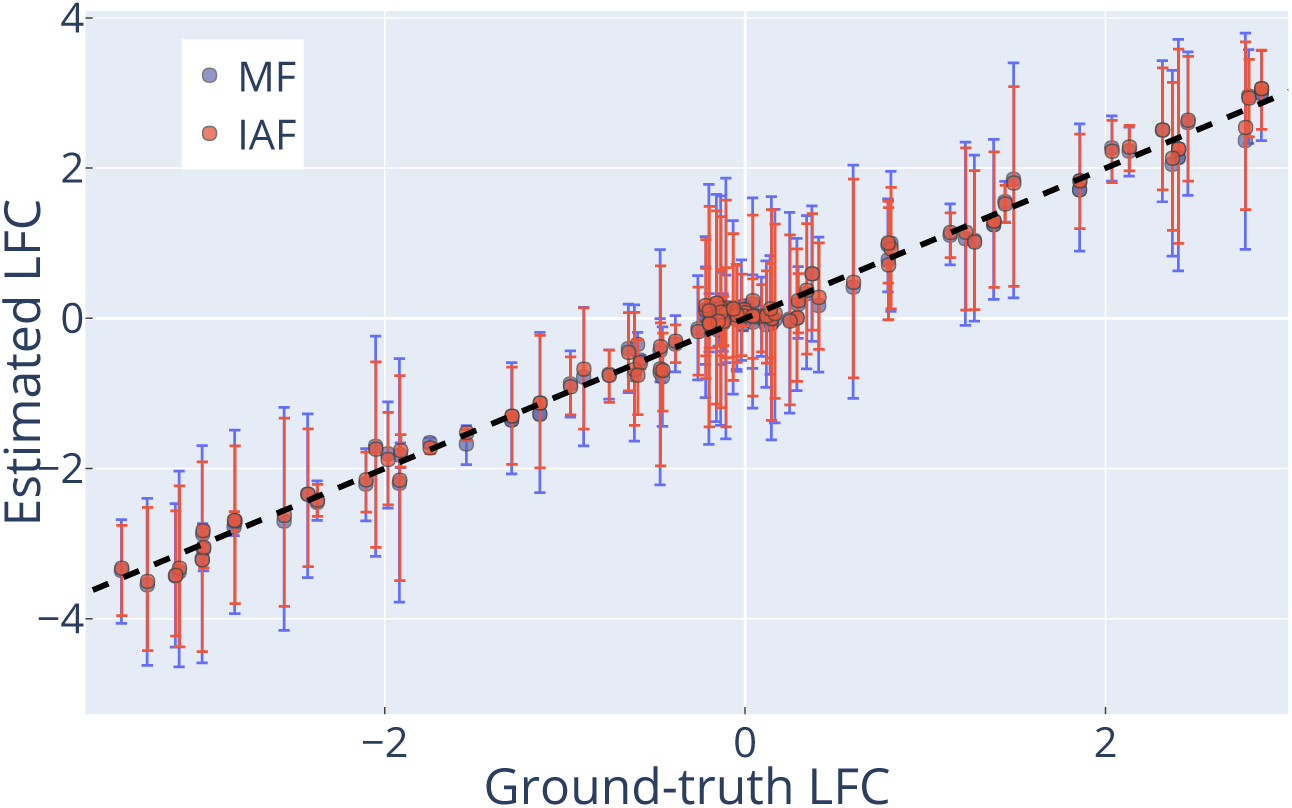
Estimated LFC against ground-truth values of the PL dataset. Points and bars respectively represent medians and 95%-credible intervals.

The results above do not assure that our LFC estimation is robust to scRNA-seq inherent biases, which can be found inn real datasets. Even though ground-truth LFC values are usually intractable on real data, LFCs estimates from scRNA-seq real datasets can be compared to reference LFC estimates computed on bulk data [18, 19], which suffers less from technical artifacts. scVI estimates are closer to the reference than the other algorithms by a large margin (Table 2), with best result achieved by IAF. Similarly, scVI best estimated the LFC for a second pair of cell-types (CD4 and CD8 cells, results not shown). This suggests that the observation model of scVI yields a reasonable choice of technical-noise free gene expression level *h*_*ng*_. Additionally, we verified (as in [8]) that our LFC estimates between subsets of cells of the same type were low (results not shown).

**Table 2:**
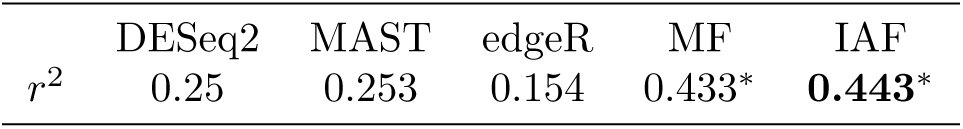
*r*^2^ coefficients of the linear regression of microarray LFCs on estimated LFCs between B cells and dendritic cells of the PBMC dataset.

|  | DESeq2 | MAST | edgeR | MF | IAF |
| --- | --- | --- | --- | --- | --- |
| $r^2$ | 0.25 | 0.253 | 0.154 | 0.433* | **0.443\*** |

### 3.3 Detection of differentially expressed genes

This section investigates the properties and use-cases of the differential expression probabilities 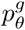 computed using MF and IAF compared to the p-values provided by competing algorithms.

We first benchmark the different algorithms on the PL dataset for which differentially expressed genes are known. In the precision-recall (PR) curves of Figure 2, MF and IAF clearly outperform other algorithms in terms of average precision and overall classification performance. Furthermore, our approach offers the best PR tradeoffs.

**Figure 2:**
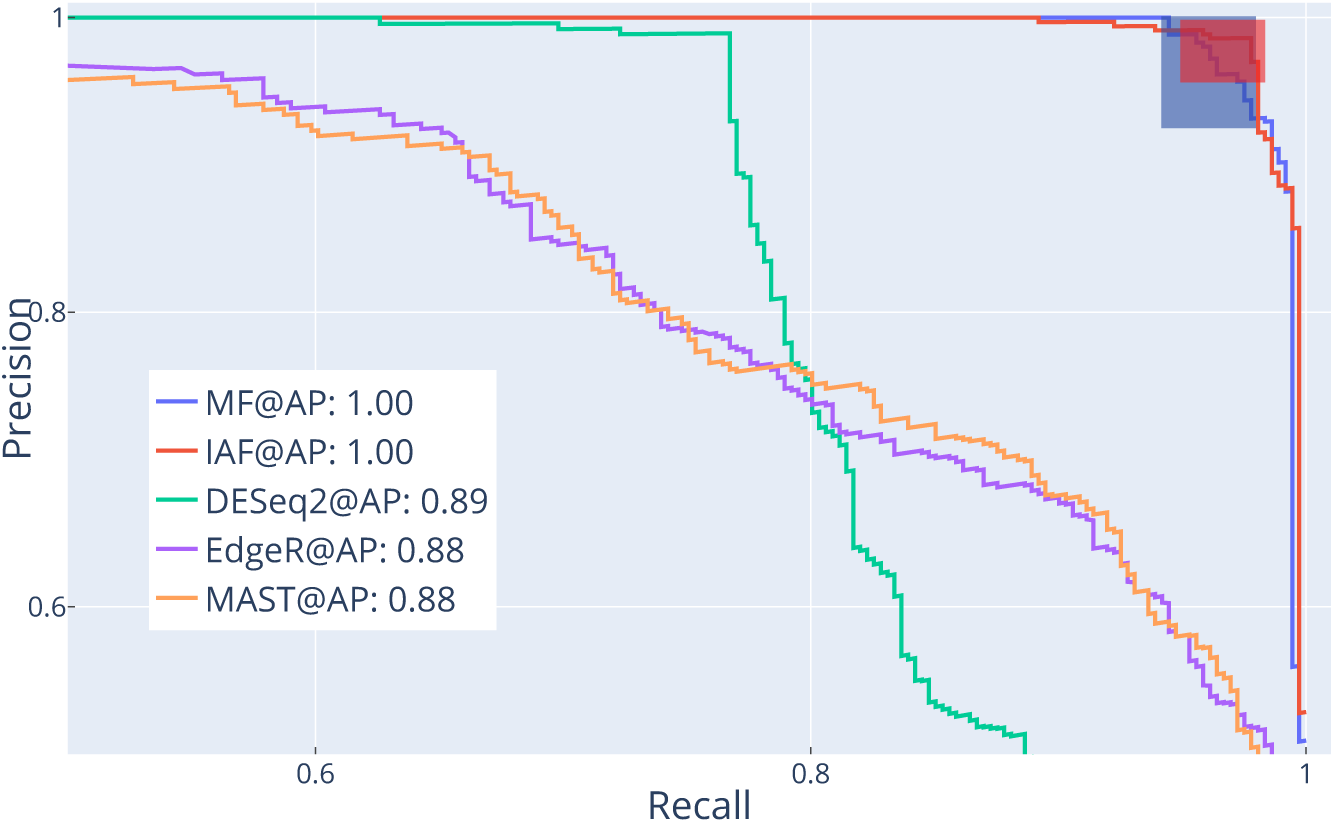
PR curves on the PL dataset for queries of size 100. Prediction scores are based on Equation 1 for IAF and MF and on significance values for other algorithms. The rectangles correspond to the 95%-confidence regions (across samplings and weight initializations) of the scVI decision rules (*α* = 0.5) characteristics.

However, those results do not assure the relevance of our protocol to detect DE genes. The two color areas on the figure correspond to the decision rule characteristics of both posteriors. IAF decision rule looks better calibrated than MF, as it provides slightly better recalls, and higher, more stable precisions (statistically significant). By better modelling uncertainty, IAF is hence able to detect more DE genes.

Differential expression techniques often require additional filtering steps after a significance test to detect differentially expressed genes of biological interest. Our DE definition takes both significance and effect size considerations into account, allowing to detect pertinent genes. Table 3 displays the ratio of predicted DE genes that have low groundtruth LFC (thus representing cases of likely little biological interest). DESeq2, followed by IAF and MF offer the best performance. The gap between DESeq2 and our approach can be explained by the fact that DESeq2 has a very conservative decision rule, and that it also formulates a composite null hypothesis. This table also shows that a complex variational distribution can help detect more DE genes of interest.

**Table 3:** Mean ratio of predicted DE genes whose LFC magnitude is lower than *δ* on the PL dataset.

| DESeq2 | MAST | edgeR | MF | IAF |
| --- | --- | --- | --- | --- |
| $3.10^{-4}$ | 0.206 | 0.042 | 0.013 | **0.008** |

The quality of the differential expression probabilities 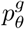 can also be assessed via SymSim, a more challenging simulation framework that explicitly models the scRNA-seq gene counts measurement process. While it is not straightforward to obtain a list of differentially expressed genes from this simulation, the LFCs can be estimated from the simulation’s model and be used as a measure of the degree of differential expression. We thus consider the genes ranking task instead of differential expression predictions to benchmark the different algorithms differential expression scores. We rank genes based on estimated DE probability for scVI-based protocols and p-values for the other algorithms. Our protocol provides significantly more informative ranking scores than other techniques on a challenging synthetic dataset (Table 4). More expressive posteriors like IAF provide more accurate rankings than mean-field. This suggests that normalizing flows helps learning a more robust and accurate generative model *p*_*θ*_.

**Table 4:** Means of Spearman correlation between DE scores and estimated LFC amplitude across experiments on Symsim.

| DESeq2 | MAST | edgeR | MF | IAF |
| --- | --- | --- | --- | --- |
| 0.428 | 0.568 | 0.032 | 0.669* | **0.681\*** |

The quality of our differential expression detection can also be assessed on real data. More precisely, the properties of DE predictions on the PBMC dataset are presented in Figure 3. We observe that IAF predictions match most of its competitors, but also include more genes hinting that it might offer better recall on real data.

**Figure 3:**
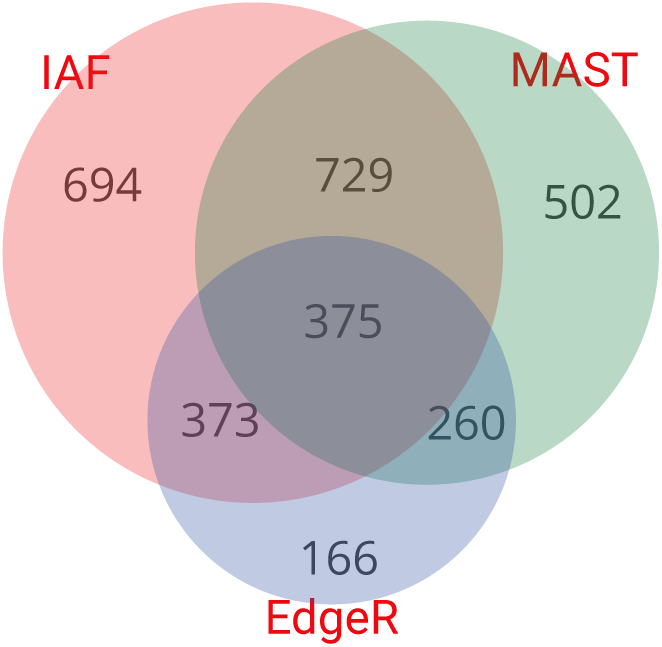
Venn diagrams of predicted DE genes (IAF, MAST, edgeR) on PBMC.

In addition, proper uncertainty quantification of IAF for DE prediction can be observed on real data. We study the number of genes incorrectly detected as DE on the PBMC dataset by taking pairs of 100 samples from the B cell population, repeated 10 times. Based on the described pipeline, IAF has only one false positive on average.

We presented a framework tailored for deep generative models for differential expression. Thanks to its data-adaptive nature and non-linearity assumptions, it successfully manages to provide reliable, informative LFC estimates, in addition to pertinent DE gene candidates for scRNA-seq data. Notably, such a framework is flexible and can be extended to other types of hypotheses such as differential variance analysis [20, 21].

## Code availability

The implementation to reproduce the experiments of this paper is available at https://github.com/PierreBoyeau/lfc_estimation. The reference implementation of scVI is available at https://github.com/YosefLab/scVI.

## References

[1] Allon Wagner, Aviv Regev, and Nir Yosef. Revealing the vectors of cellular identity with single-cell genomics. Nature biotechnology, 2016.

[2] Alessandra Dal Molin, Giacomo Baruzzo, and Barbara Di Camillo. Single-Cell RNA-Sequencing: Assessment of Differential Expression Analysis Methods. Frontiers in Genetics, 2017.

[3] Greg Finak, Andrew McDavid, Masanao Yajima, Jingyuan Deng, Vivian Gersuk, Alex K. Shalek, Chloe K. Slichter, Hannah W. Miller, M. Juliana McElrath, Martin Prlic, Peter S. Linsley, and Raphael Gottardo. MAST: a flexible statistical framework for assessing transcriptional changes and characterizing heterogeneity in single-cell RNA sequencing data. Genome Biology, 2015.

[4] Peter V Kharchenko, Lev Silberstein, and David T Scadden. Bayesian approach to single-cell differential expression analysis. Nature Methods, 2014.

[5] Rahul Satija, Jeffrey A. Farrell, David Gennert, Alexander F. Schier, and Aviv Regev. Spatial reconstruction of single-cell gene expression data. Nature Biotechnology, 2015.

[6] Michael I Love, Wolfgang Huber, and Simon Anders. Moderated estimation of fold change and dispersion for RNA-seq data with DESeq2. Genome Biology, 2014.

[7] Mark D. Robinson, Davis J. McCarthy, and Gordon K. Smyth. edgeR: a Bioconductor package for differential expression analysis of digital gene expression data. Bioinformatics, 2010.

[8] Charlotte Soneson and Mark D. Robinson. Bias, robustness and scalability in single-cell differential expression analysis. Nature Methods, 2018.

[9] Romain Lopez, Jeffrey Regier, Michael B. Cole, Michael I. Jordan, and Nir Yosef. Deep generative modeling for single-cell transcriptomics. Nature Methods, 2018.

[10] James Berger. Statistical Decision Theory and Bayesian Analysis. Second edition, 1985.

[11] Shiqi Cui, Subharup Guha, Marco A. R. Ferreira, and Allison N. Tegge. hmmSeq: A hidden Markov model for detecting differentially expressed genes from RNA-seq data. The Annals of Applied Statistics, 2015.

[12] Diederik P. Kingma and Max Welling. Auto-Encoding Variational Bayes. In International Conference on Learning Representations, 2014.

[13] Richard E Turner, Pietro Berkes, and Maneesh Sahani. Two problems with variational expectation maximisation for time-series models. Inference and Estimation in Probabilistic Time-Series Models, 2010.

[14] Diederik P. Kingma, Tim Salimans, Rafal Jozefowicz, Xi Chen, Ilya Sutskever, and Max Welling. Improving Variational Inference with Inverse Autoregressive Flow. In Neural Information Processing Systems, 2016.

[15] Charlotte Soneson and Mark D. Robinson. Bias, robustness and scalability in single-cell differential expression analysis. Nature Methods, 2018.

[16] Xiuwei Zhang, Chenling Xu, and Nir Yosef. SymSim: simulating multi-faceted variability in single cell RNA sequencing. Nature Communications, 2019.

[17] Volodymyr Kuleshov, Nathan Fenner, and Stefano Ermon. Accurate uncertainties for deep learning using calibrated regression. International Conference on Machine Learning, 2018.

[18] Helder I. Nakaya, Jens Wrammert, et al. Systems biology of vaccination for seasonal influenza in humans. Nature Immunology, 2011.

[19] Holderried Tobias Görgün, Güllü et al. Chronic lymphocytic leukemia cells induce changes in gene expression of CD4 and CD8 T cells. The Journal of clinical investigation, 2005.

[20] Zhun Miao, Ke Deng, Xiaowo Wang, and Xuegong Zhang. Desingle for detecting three types of differential expression in single-cell rna-seq data. Bioinformatics, 2018.

[21] Nils Eling, Arianne C Richard, Sylvia Richardson, John C Marioni, and Catalina A Vallejos. Correcting the mean-variance dependency for differential variability testing using single-cell rna sequencing data. Cell systems, 2018.

